# Single-cell phenotyping of human induced pluripotent stem cells by high-throughput imaging

**DOI:** 10.1101/026955

**Authors:** H. Christian Volz, Florian Heigwer, Tatjana Wüest, Marta Galach, Jochen Utikal, Hugo A. Katus, Michael Boutros

**Affiliations:** German Cancer Research Center (DKFZ), Division Signaling and Functional Genomics and Heidelberg University, Heidelberg, Germany; Department of Cardiology, University Hospital Heidelberg, Heidelberg, Germany; German Cancer Research Center (DKFZ), Skin Cancer Unit, Heidelberg and Department of Dermatology, Venereology and Allergology, University Medical Center Mannheim, Heidelberg University, Mannheim, Germany; DZHK (German Centre for Cardiovascular Research), partner site Heidelberg/Mannheim, Germany

**Keywords:** stem cells, automated imaging, single-cell phenotyping, bioinformatics

## Abstract

Single-cell phenotyping promises to yield insights into biological responses in heterogeneous cell populations. We developed a method based on single-cell analysis to phenotype human induced pluripotent stem cells (hIPSC) by high-throughput imaging. Our method uses markers for morphology and pluripotency as well as social features to characterize perturbations using a meta-phenotype based on mapping single cells to distinct phenotypic classes. Analysis of perturbations on a single cell level enhances the applicability of human induced pluripotent stem cells (hIPSC) for screening experiments taking the inherently increased phenotypic variability of these cells into account. We adapted miniaturized culture conditions to allow for the utilization of hIPSC in RNA interference (RNAi) high-throughput screens and single cell phenotyping by image analysis. We identified key regulators of pluripotency in hIPSC masked in a population-averaged analysis and we confirmed several candidate genes (*SMG1*, *TAF1*) and assessed their effect on pluripotency.

## Introduction

Multi-parametric analysis applied to high throughput imaging screens opened new avenues in functional genomic analysis.^1^ While this method allows to measure the effect of perturbations by small molecules or small interfering RNAs (siRNAs) with cellular resolution^2,3^, robust methods are necessary to analyse the vast amounts of multidimensional data on a single-cell level.^4,5^ Currently, most methods reduce data complexity by normalization and averaging over cell populations, thereby loosing information about phenotypic plasticity and heterogeneity within a population of perturbed cells.^6,7^ In contrast, preserving the multi-dimensionality at a single-cell level captures cell-to-cell variability and promises to provide in-depth characterization of heterogenic cell populations, such as stem cells, that are otherwise not achievable. Such analysis can be provided by multi-factorial microscopy based readout.

Human induced pluripotent stem cells (hIPSC) are an important model system to address fundamental questions in stem cell and developmental biology^8–10^, and serve as a tool to dissect disease processes using patient-derived hIPSC. The phenotypic heterogeneity of hIPSC, in particular after perturbations that might interfere with self-renewal or initiate differentiation, requires a quantitative, single-cell analysis of hIPSCs.^11–13^ Here we established a high-throughput method to comprehensively measure multi factorial phenotypes in hIPSC perturbed by RNA interference (RNAi) in order to overcome limitations for hIPSC use in high-throughput screening applications.

## Results

To identify modifiers of hIPSC phenotypes, we performed a kinome-wide RNAi screen using automated microscopy (Figure 1a,b). First, we established a method to reliably seed, perturb and image hIPSC in a 384-microwell format. Transparent-bottom micro well plates were coated with Matrigel and siRNAs were pre-spotted. After treatment with ROCK-inhibitor Y27632 for 1h followed by enzymatic digestion with trypsin and collagenase IV, 2000 separated hIPSC per well were seeded and reverse transfected with siRNAs in mTeSR™ stem cell medium. We found conditions to ensure that cells were evenly distributed, spatially separated and growing robustly (Figure 2a). After 4 days incubation, cells were fixed and stained for DNA and OCT4 protein expression. Next, we tested the gene silencing efficiency of small interfering RNAs (siRNA) targeting *OCT4* (*Pou5f1*), a master regulator of pluripotency in hIPSC, marking cellular pluripotency state.

**Figure 1.**
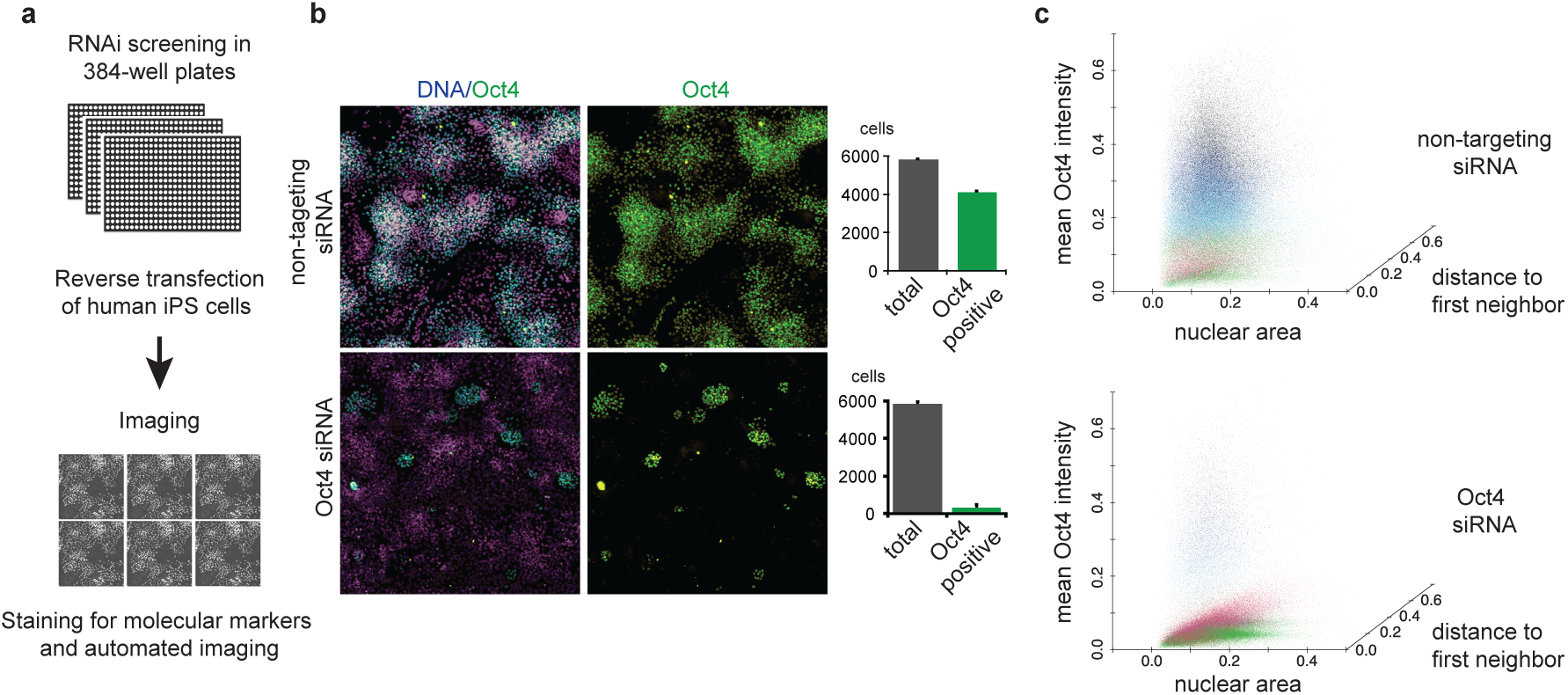
Single-cell phenotyping of human induced pluripotent stem cells. (a) Workflow of cell-based RNAi screening of hIPSC using a high-throughput microscopy read-out. (b) Representative images of hIPSC 96 hours after transfection with siRNA targeting Oct4 and non-targeting control siRNA. Bar plots show quantification of Oct4-positive and total nuclei. (c) Distinct populations of hIPSC. Each cell is represented by a dot with the following feature data: nuclear area, mean Oct4 signal intensity, and distance to first neighbour. 200 000 randomly selected cells from both, non-targeting and Oct4 targeting siRNA controls were plotted. Feature data were scaled between 0 and 1 linearly, based on the complete dataset. Colours represent populations found by k-means clustering with 6 centres and 30 random starts.

A pool of 4 siRNAs targeting *OCT4* efficiently silenced OCT4 expression reducing the number of OCT4 positive cells 13-fold (Figure 1b). Additional markers of pluripotency, SSEA4 and NANOG decreased 80 fold and 6 fold after OCT4 knockdown, respectively (Figure 4b). Subsequently, a total of 1.71 Mio cells were seeded in two biological replicates. Automated imaging was performed using a robotic imaging system to acquire more than 42,000 images. We developed an image analysis pipeline for hIPSC analysis based on CellProfiler^14^ (Suppl. Software 1). Quantified features included intensity, morphology and spatial distribution features (see also Methods, Image analysis). An analysis on the basis of the z’-score normalized OCT4 signal showed overall good reproducibility of the biological replicates (Figure 2b, Spearman correlation coefficient of 0.67). In-depth analysis of the screen based on multiple single-cell features was realized using a workflow based on single-cell feature clustering approaches (Figure 2d, Suppl. Data 1).^7^ Out of an initial dataset of 67 features, non-redundant features were selected by excluding features showing Pearson correlation coefficients of higher than 0.95 with any other feature. For subsequent analysis, we used 8 remaining features: (i) nuclear extent, (ii) nuclear area, (iii) number of neighbours, (iv) distance to first neighbour, (v) mean Hoechst intensity, (vi) maximum Hoechst intensity, (vii) mean OCT4 intensity and (viii) maximum OCT4 intensity. Unsupervised k-means clustering was performed on negative (non-targeting siRNA) and positive (siRNA targeting *OCT4*) control cells resulting in a classification into multiple distinct cell categories.

**Figure 2.**
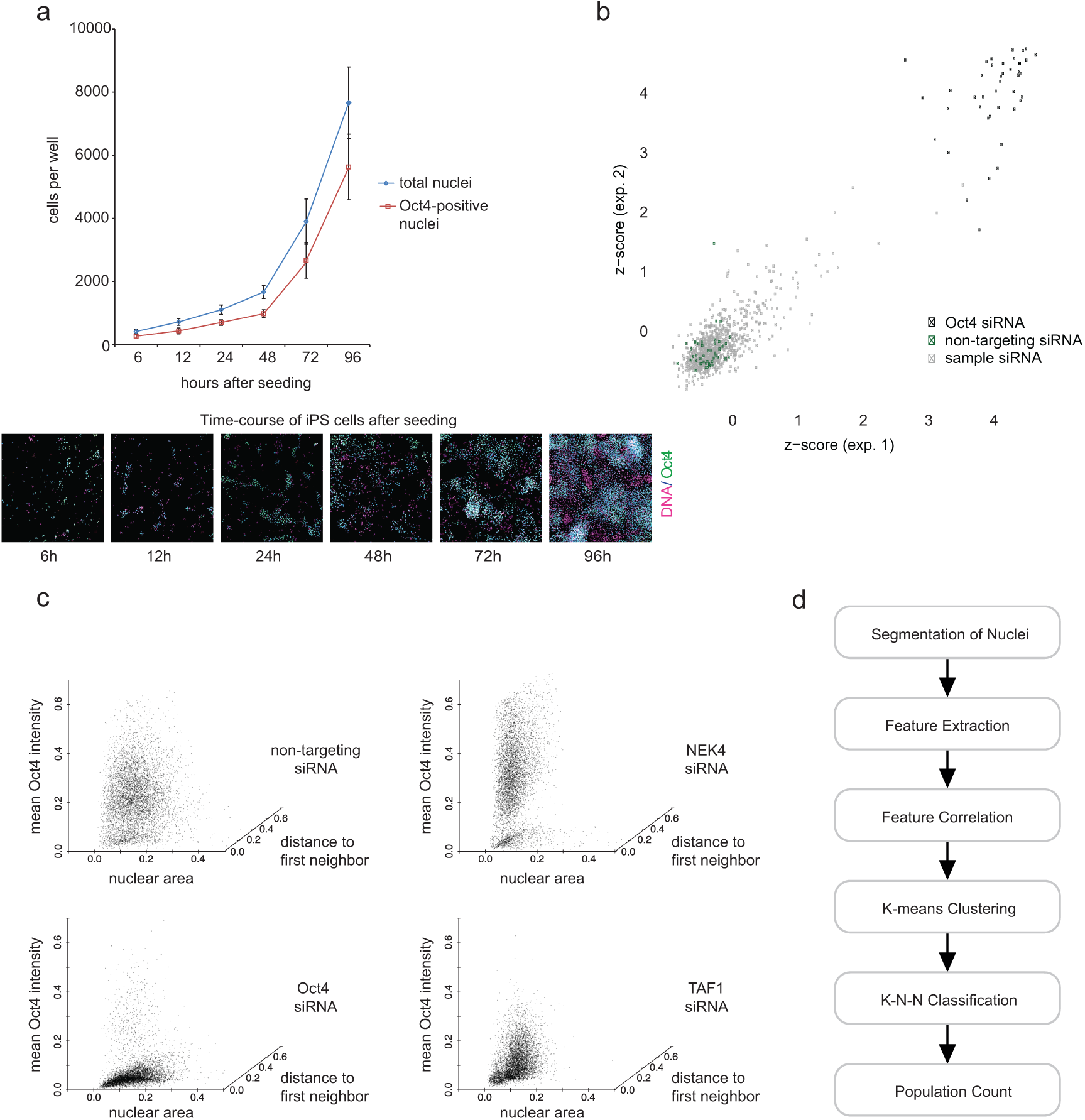
Methods for single cell phenotyping in hIPSC. (a) Growth curve of hIPSC after seeding 2000 single cells per well in a 384-microwell plate (n = 8 per time point, mean ± s.d.). Corresponding immunofluorescence images of hIPSC stained for DNA (purple) and Oct4 (green) after single-cell seeding. (b) Scatterplot of two replicate RNAi screening results using per-well analysis of 779 sample siRNAs targeting kinases (grey) and 21 positive (siRNA targeting OCT4; black) as well as 42 negative (non-targeting siRNAs; green) control siRNAs, respectively. Z-scores were calculated based on the ratio of the number of Oct4-positve cells and the total cell number per siRNA-treatment (Spearmen correlation coefficient = 0.67). (c) The three features (I) nuclear area, (II) mean Oct4 signal intensity, and (III) distance to first neighbour were plotted for non-targeting siRNA, *OCT4*-siRNA, *NEK4*-siRNA and *TAF1*-siRNA, respectively. The cells were subsampled from the entire dataset of each treatment, randomly 6000 cells from each treatment. The data was scaled linearly from 0 to 1. (d) Step-by-step schematic workflow comprising image segmentation, feature extraction, reduction of the feature matrix by correlation analyses, K-means clustering to generate phenotypic population categories, K-nearest-neighbour (K-N-N) classification by assigning every single cell of the screening experiment to one of the populations, and the final profiling by counting the number of cells of each population in every well.

Classification into six categories resulted in the most robust clustering performance. We visualized these six reference cell categories in a 3D scatterplot (i) nuclear area, (ii) OCT4-signal intensity, and (ii) distance to first neighbour after treatment with non-targeting siRNA and siRNA targeting *OCT4*. OCT4 signal intensity was reduced by 80 % after *OCT4* knockdown. In addition, reduction in the expression of the remaining two features could also be detected, 3.5 % and 2 % for nuclear area and distance to first neighbour, respectively (Figure 2c, left). This highlights the shift of entire cell populations after down regulation (Figure 1c, e.g. reduction of abundance of ‘black’ category cells by nearly 100 %, see also Figure 3b class 5). Reference categories were further used to perform k-nearest neighbour classification assigning each cell of the 7731725 imaged cells to one of the six categories (Figure 3a). We further found that the population distribution of the cells among the reference categories can be used as a characteristic meta-phenotype and to enable the classification of perturbations. Six profile distributions, showing enrichments for either one population, are shown in Figure 3b, including example perturbations. For example, knockdown of NIMA-related kinase 4 (*NEK4*) increases the non-targeting (stem cell-like) meta-phenotype, whereas in contrast RNAi against the TATA box binding protein (TBP)-associated factor (*TAF1*) is forcing the population into an *OCT4*-knockdown-like distribution (Figure 2c).

**Figure 3.**
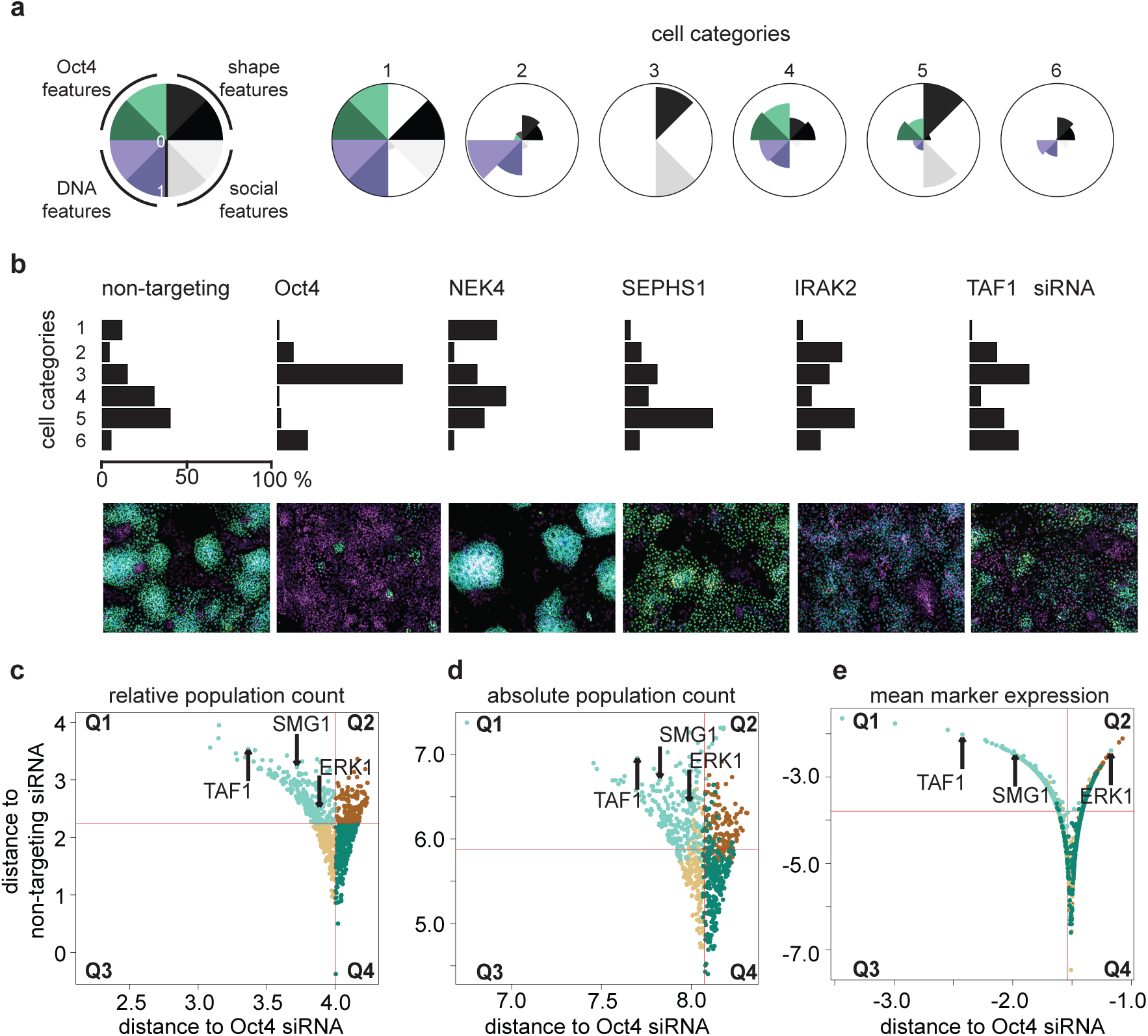
Profiling by single-cell classification and profile correlation. (a) 6 categories defined by k-means clustering are represented by star plots of the 8 phenotypic traits used. The following traits were represented: number of neighbours (light grey), distance to first neighbour (dark grey), mean Hoechst intensity (light purple), maximum Hoechst intensity (dark purple), mean Oct4 intensity (dark green), maximum Oct4 intensity (light green), nuclear extent (dark grey), nuclear area (black). (b) Population profiles of selected transcript depletions. For each profile, cells in each population were counted and normalized to the total cell count for the corresponding treatment. Bar plots represent the distribution of cells among the 6 categories. In addition an image enriched for either one population is shown beneath emphasizing the visual phenotype of the 6 populations. (c-e) Euclidean distance of each sample profile from the average non-targeting siRNA profile is plotted against the Euclidean distance of each sample profile from the average Oct4 targeting siRNA controls. (c) Profiles are defined as the per cent cell counts of each population normalized to the total number of cells per well. (d) Profiles are defined as absolute cell counts per population per well. (e) Profiles are defined as mean feature expression per well regarding mean and maximum Oct4 signal intensity.

Perturbations are characterized based on the Euclidian distance of their meta-phenotype to positive and negative controls. When plotted against two controls, candidate genes were segregated into four groups (Figure 3c-e), showing e.g. either close proximity to positive controls (Q1) and far proximity to negative controls or far proximity to positive and negative controls (Q2). For instance, perturbations resulting in a shift of a profile from Q3 to Q1 represent the transformation from a pluripotent meta-phenotype towards a loss-of-pluripotency. In contrast, meta-phenotypes located in Q2, differ from an unperturbed hIPSC phenotype, however without resembling the loss-of-pluripotency meta-phenotype. For example, depletion of *TAF1* or *SMG1* gene in hIPSC shows a meta-phenotype located in Q1 indicating close phenotypic relation to the *OCT4*-knockdown (Figure 3d). However, whereas the meta-phenotype of *ERK1*, a known regulator of pluirpotency^15^, shows similarities to the *OCT4* control (Q1), an analysis of OCT4 expression only (Figure 3e) does not implicate this and locates the meta-phenotype in Q2. Even if cell count per population is normalized to total number of cells, excluding viability effects, the same trends are observed (Figure 3d).

We further validated the two candidate genes, *TAF1* and *SMG1*, using deconvoluted pools of siRNAs. In addition, we analysed expression of additional pluripotency markers other than Oct4 after targeting *TAF1* and *SMG1* with siRNA. These experiments demonstrated effective gene silencing of *TAF1 and SMG1* with RNAi and revealed decreased expression of NANOG and SSEA4 indicating loss of pluripotency in hIPSC after knockdown of *TAF1* and *SMG1* (Figure 4c).

**Figure 4.**
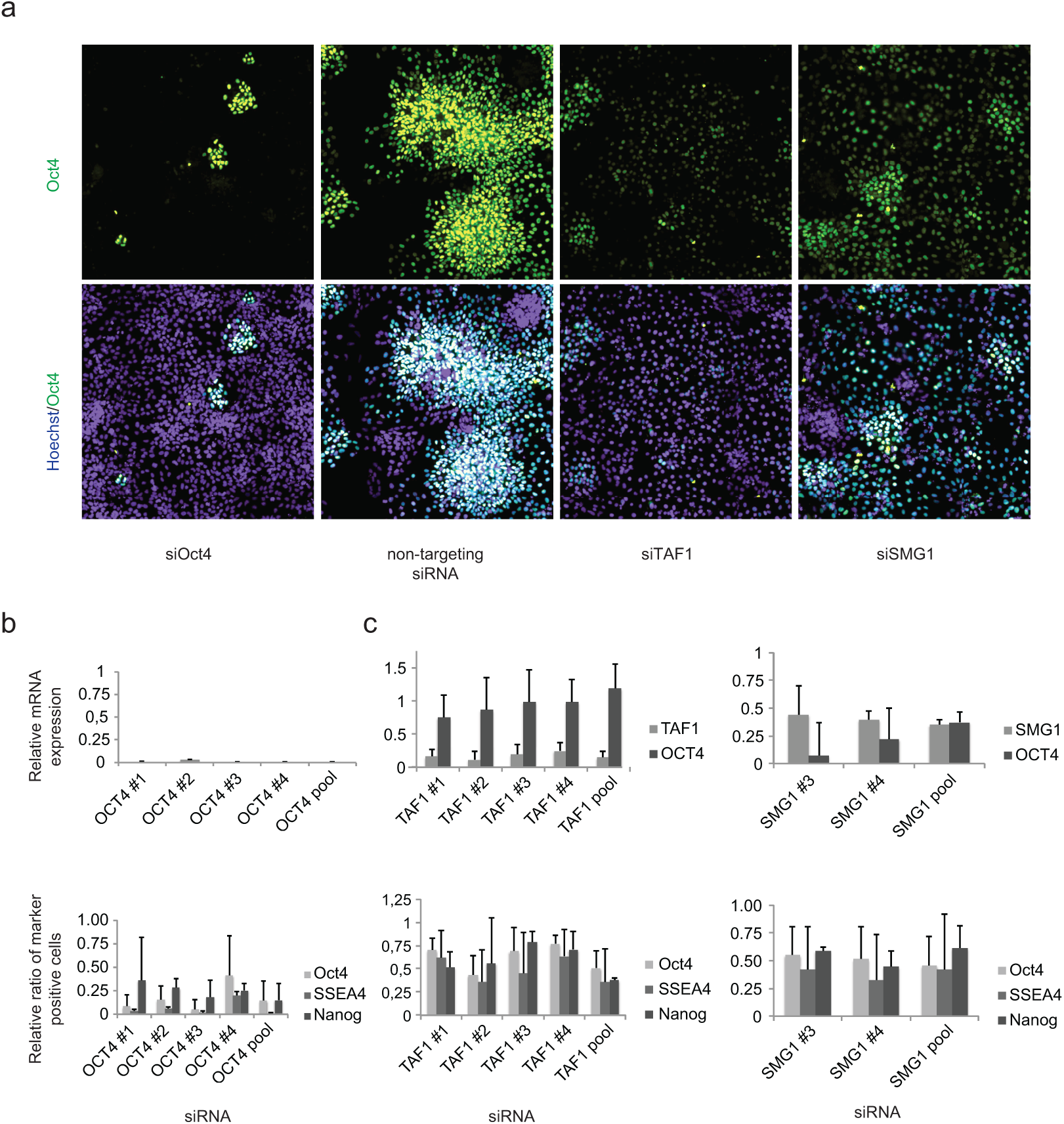
Follow–up analysis of the candidate genes TAF1 and SMG1. (a) Immunofluorescence images of hIPSC stained for DNA (blue) and Oct4 (green) after treatment with siRNAs targeting *OCT4*, *TAF1*, *SMG1*, and with non-targeting siRNA. (b) *OCT4* gene expression, measured by qRT-PCR and normalized to *ACTB* mRNA levels, and relative ratio of OCT4-, SSEA4- and NANOG-positive hIPSC transfected with single siRNAs and corresponding pools of siRNAs targeting *OCT4*. Data shown as mean ± s. d. (n = 5 for gene expression data; n = 4 for OCT4, n = 2 for SSEA4 and NANOG immunofluorescence data). (c) Expression of candidate genes *TAF1* and *SMG1,* and of pluripotency marker *OCT4* normalized to *ACBT* in hIPSC transfected with single siRNAs and the corresponding pool of siRNAs targeting *TAF1* und *SMG1*, respectively. Data shown as mean ± s. d. (*TAF1*-knockdown: n = 3 for *TAF1* and n = 5 for *OCT4*; *SMG1*-knockdown: n = 2 for *SMG1* and n = 5 for *OCT4*). Relative ratio of cells positive for pluripotency markers OCT4, SSEA, and NANOG in hIPSC transfected with single siRNAs and the corresponding pool of siRNAs targeting *TAF1* und *SMG1*, respectively. Data shown as mean ± s. d. (n = 2 – 4 for *TAF1*, n = 2 – 4 for *SMG1*).

## Conclusions

Large-scale functional analyses of heterogeneous cells have remained an unresolved challenge for high-throughput experiments. Our combined experimental and computational workflow demonstrates how image-based single-cell phenotyping could be used to identify candidate perturbations. We show how perturbation of these gene impact phenotypic characteristics of cell populations.

This indicates that cell populations undergo different functional states such as the modulation of the pluripotent state as shown in our study. Combining morphology-based features with specific markers robustly identified phenotypic classes for a heterogeneous cell type at a single-cell level.^16^ Future experiments using additional markers or different control profiles may aid in the assessment of differentiation capacity of hIPSC towards specific phenotypes and help to identify perturbations that influence specific differentiation pathways.

## Methods

### Cell Culture

Experiments were performed in two human induced pluripotent stem cell (hIPSC) lines (hIPSC1 and hIPSC2, as indicated in Suppl. Table 1). hIPSC1 was derived from human skin fibroblasts (Ethics Committee of Heidelberg University approval no. 2009-350N-MA) and hIPSC2 was generated using replication-defective doxycycline-inducible single lentiviral vectors containing OCT4, KLF4 and C-MYC from primary melanocytes purchased from Promocell as described previously^1^. Reprogramming of hIPSC1 was performed using the polycistronic lentiviral vector STEMCCA-OKSM and the tet-activator *M2rtTA* as described elsewhere^2^. Fibroblasts were seeded with a density of 5000 cells/well on gelatin coated 6-well plates (BD Bioscience). The cells were rinsed with PBS and 2 mL mouse embryonic fibroblast (MEF) medium (DMEM supplemented with 10% fetal bovine serum, 1% penicillin/streptomycin and 1% L-glutamine, all Gibco) was added. For lentiviral transduction 5 μL of M2rtTA and 5 μL STEMCCA-OKSM concentrated virus was added. To improve the transduction efficiency 1 mg/mL polybrene (Sigma) was supplemented. The following day cells were rinsed with PBS followed by another round of lentiviral transduction. After 24 h the cells were washed twice with PBS and a 1:1 mixture of MEF medium with human embryonic stem cell (hES) medium (DMEM/F12 supplemented with 20 % KOSR, 1% L-glutamine, 1% penicilline/streptomycine, 1% MEM-NEAA, all Gibco, 0.05 mM 2-Mercaptoethanol, Carl Roth, and 10 ng/mL bFGF, Promokine) was added. The medium was supplemented with 1 μg/mL doxycycline (Sigma) in order to induce the transgene expression. The transduced cells were cultured in the mix-medium for one week, afterwards in hES cell medium supplemented with 1 μg/mL doxycycline. After four to six weeks appearing colonies of reprogrammed cells were picked manually and transferred onto mitomycin c (AppliChem) treated feeder cells. After another two weeks, when the cells reached a stable state, doxycycline was withdrawn and the colonies were picked manually onto Matrigel™ (BD Bioscience) coated dishes and cultured in mTeSR™ (STEMCELL Technologies). To ensure pluripotency hIPSC1 were subjected to a teratoma formation assay and the expression of pluripotency markers was assessed by immunostaining (Suppl. Figure 1).

Both, hIPSC1 and hIPSC2 were cultured in mTeSR™1 (STEMCELL Technologies) in feeder-free conditions on growth factor reduced Matrigel-coated (BD Bioscience) plates in a humidified incubator at 37° C and 5% CO_2_ in standard conditions.

### High-throughput siRNA screening

The kinome siRNA library (siGENOME, Dharmacon) was arrayed in growth factor reduced Matrigel (BD Bioscience) coated (20 μL per well for one hour at room temperature) 384 well micro plates (BD Bioscience) using a Biomek FX200 liquid handling system. 5 μL of a 250 nmolar siRNA pool of four single siRNAs were spotted in each well. Non-targeting single siRNA (siControl #2 in wells B4 through H4 and siRLUC in wells I4 through O4, Dharmacon) served as negative and a pool of four siRNAs targeting OCT4 (SiGenome, Dharmacon; wells B3 through H3) served as positive controls. A pool of four single siRNAs targeting UBC (SiGenome, Dharmacon; wells A3, A4, P3, and P4) served as viability controls.

For single cell seeding hIPSC were treated with 5 μmolar ROCK-inhibitor (Y27632; Axxora, 5 μmolar)) in mTeSR™1 hour before and after passaging. Cells were passaged in a two-step protocol sequentially using CTK solution consisting of 0.25% trypsin (Gibco), 0.1% collagenase IV (Invitrogen), 20% KnockOut Serum Replacement (Invitrogen) and 1 mmolar CaCl_2_ as well as trypsin solution (0.25% with EDTA; Gibco) to generate a single cell suspension. Reverse transfection was performed using Lipofectamine RNAiMAX (Invitrogen) transfection reagent. A master mix of 0.05 μL transfection reagent and 4.95 μL DMEM/F12 (Invitrogen) medium was added to each well and incubated for 20 minutes. Single hIPSC were seeded at a density of 2000 cells per well in 40μL mTeSR™1 medium supplemented with 1% penicillin/streptomycin (Invitrogen) and 5 μmolar ROCK-inhibitor. Two days after transfection 30 μL of fresh mTeSR™1 was added to each well. Plates were kept in a humidified incubator at 37° C and 5% CO_2_ in standard conditions.

### Fixation and Staining

Four days after transfection, cells were washed with PBS. Subsequently cells were fixed and permeabilized using 4% paraformaldehyde (Sigma) and 0.2% Triton X-100 (AppliChem) in PBS for 45 minutes at room temperature. After an additional washing step with PBS, cells were blocked with 3% BSA (Sigma) and 0.05% Triton X-100 in PBS for 45 minutes. Next, immunostaining was performed overnight at 4° C using a rabbit anti-OCT4 antibody (Abcam, working concentration 1 μg/mL) and Phalloidin-TRITC (Sigma; working concentration 1 μg/mL). After three washing steps secondary antibody staining was performed with an Alexa 488 conjugated goat anti-rabbit antibody (Invitrogen; working concentration 5μg/mL) and DNA-labelling was done with Hoechst 33242 (Invitrogen; working concentration 1μg/mL) for 1 hour at room temperature. After 4 additional washing steps plates were amenable for image acquisition.

### Automated imaging

Imaging was done using an automated BD Pathway 855 Bioimaging System (Becton Dickinson) with a 20x objective (NA=0.75) in combination with a Hamamatsu digital camera (Orca-ER). 25 fields per well were imaged containing channels for Hoechst (DNA; excitation filter 380/10 nm), and FITC (Oct4; excitation filter 488/10 nm). Hoechst staining was used to bring cells into focus by laser autofocus for every single image. Due to a technical restraint of the BD Pathway microscope leading to partial duplication of image frames in row A of 384 micro well plates, all compound images had to be cropped accordingly to prevent duplicate frames to be analysed.

### Image analysis

Image analysis was performed with the CellProfiler image analysis software (Version 2.0). The pipeline used in this screen is available under Suppl. Software 2. In brief, object selection was based on adaptive intensity and fixed size thresholds for every single object. Object segmentation was optimized to achieve best possible resolution of single objects in dense clusters of cells. For data analysis, morphological, intensity and spatial distribution parameters of parent objects segmented in channel 1 (Hoechst) and child objects of objects in channel 2 (Oct4) were measured.

### Per-well screen analysis using cellHTS2

After manually setting a threshold for mean intensity of the Oct4 signal, cells were binned into Oct4-positive and Oct4-negative cells. The ratio of Oct4-positive nuclei (r_p_) was calculated by dividing the number of Oct4-positive objects by the total number of nuclei per well. To exclude siRNAs strongly affecting cell viability, a minimum threshold t_min_ for the cell number per well was implemented at the 5% quartile of the number of nuclei of all wells analysed for each pair of plates (plate 1: t_min_ = 2430, plate 2: t_min_ = 2967, plate 3: t_min_ = 3268). The screen was analysed using the R package cellHTS2^3^. Single channel data (fraction of Oct4-positive cells per well) were normalized per plate and z-scores were calculated (ranked list of z-scores see Suppl. Data 2). The Spearman rank correlation coefficient of the two technical duplicates of the screen was 0.67 for r_p_. The z’-factor of the ratio of Oct4-positive nuclei per well was 0.62 and 0.42 for replicate 1 and replicate 2, respectively.

### Single cell phenotyping

R was used in version 3.0.2. Perl was used in version 5.16.2. Raw data has been acquired from tab delimited output files from CellProfiler and annotated with the attached Perl scripts *find_ctrls.pl* and *map_rna_to_well.pl* (Suppl. Data 1,2). These scripts require a folder structure, in which for every plate there is a subfolder containing the respective tab delimited file and the annotation file. In each txt file there is one row per segmented object containing its well number of origin and its feature vector. The Annotation file hence contains the well number, well name and RNAi reagent. Further, using the R function *make_and_save_kmeans* (Suppl. Data 1,2) raw data resulting from *find_ctrls.pl* were read in, non-correlating feature (coefficient=0.95, method=Pearson) were selected and used for k-means clustering with 6 centres, 30 random start attempts and default parameters else. Clustering quality has been assessed by observing a between sum-of-squares vs. total sum-of-squares ratio of 66.5%. Following, sample cells were classified as belonging to one of the 6 phenotypic groups using the *classify_samples* function (Suppl. Data 1,2). Therein k-nearest-neighbour classification has been performed using the result of the k-means clustering as training dataset and a neighbourhood (k) of 100 data points. Hit identification has been performed by using the function *make_distance_plot*, where the Euclidian distances of each samples feature vector to the positive control vector is plotted against the negative control vector. (Suppl. Data 1,2).

### Validation Experiments

Knockdown efficiency of candidates was tested by transfecting the pools of siRNAs as well as single siRNAs (all Dharmacon; see Suppl. Table 1 for target nucleotide sequences) in hIPSC in a 96-well format. RNA from two wells of 96-well plate was isolated with the RNAeasy Mini kit (Qiagen) after 4 days of culture with conditions identical to the screen after up scaling of the respective volumes. RNA was used as template for cDNA synthesis with the Revert Aid H Minus First Strand cDNA Synthesis kit (Fermentas). Real time polymerase chain reaction was used to quantify expression of the targeted transcript using a Roche Lightcycler 480 (Roche) and the Roche Universal Probe Library (Roche). Gene expression was normalized against *ACTB*. Immunofluorescence antibody staining for Oct4, SSEA4 and Nanog (all Abcam, working concentrations 1.0, 10.0 and 0.4 μg/mL, respectively) and corresponding secondary antibodies (Alexa 488 goat anti-rabbit and Alexa 594 goat anti-mouse, Invitrogen; working concentrations of 5.0 and 8.0 μg/mL, respectively) was done using the same conditions as in the screen in a 384-well format after correction of the corresponding volumes. Imaging was done using the BD Pathway 855 Imaging system (BD) as previously described for the screening experiment. Ratio (r_p_) was calculated for each treatment using the above mentioned stem cell markers by dividing the number marker-positive cells by the total number of cells per well. For each experiment the mean of at least two wells for each treatment was calculated. To ensure comparability of data between individual experiments, the r_p_ of samples was normalized to the r_p_ of controls (non-targeting siRNA) to yield the relative ratio of marker positive cells.

## Acknowledgement

We thank D. Kranz, F. Graf, T. Sandmann and members of the Boutros and Utikal groups for helpful discussions. Research in the lab of M.B. is supported by an ERC advanced grant. This work has in part been supported the DFG SFB873 (“Stem cells in development and disease”). H.C.V. acknowledges the support by Klaus Tschira Stiftung.

## Author Contributions

H.C.V, J.U., H.A.K. and M.B. designed the study concept. H.C.V, T.W. performed the experiments. M.G. contributed to experiments. F.H. and H.C.V. analysed the data. H.C.V., F.H., and M.B. wrote the manuscript. All authors commented on the manuscript.

